# The timing of sleep spindles is modulated by the respiratory cycle in humans

**DOI:** 10.1101/2023.03.31.534952

**Authors:** Valentin Ghibaudo, Maxime Juventin, Nathalie Buonviso, Laure Peter-Derex

## Abstract

Coupling of sleep spindles with cortical slow waves and hippocampus sharp-waves ripples is crucial for sleep-related memory consolidation. Recent literature evidenced that nasal respiration modulates neural activity in large-scale brain networks. In the rodent, this respiratory drive strongly varies according to vigilance states. Particularly, during sleep, respiration promotes the coupling between hippocampal sharp-wave ripples and cortical DOWN/UP state transitions. However, no study has examined whether sleep spindles could be respiration-modulated in humans. In this work, we aimed to investigate the influence of breathing on brain oscillations during non-rapid-eye-movement stage 2 sleep (N2) in humans by examining the coupling between sleep spindles and respiration cycle. Full night polysomnography of twenty healthy participants were analysed. Spindles and slow waves were detected during N2 sleep stage. Spindle-related sigma power as well as spindle and slow waves events were analysed according to the respiratory phase. We found a significant coupling between slow and fast spindles with respiration cycle, with enhanced sigma activity and probability of occurrence of spindles during the middle part of the expiration phase. A different coupling was observed between breathing and slow waves that were more distributed around both respiration phase transitions. Our findings suggest that breathing cycle influences the dynamics of brain activity during non-rapid-eye-movement sleep. This may enable sleep spindles to synchronize with other brain rhythms including hippocampus sharp wave ripples and facilitate information transfer between distributed brain networks.

## INTRODUCTION

Sleep spindles are brief (typically 0.5-2 s) bursts of sigma (10-16 Hz) oscillations recorded in the electroencephalogram (EEG) during non-rapid-eye-movement (NREM) sleep. They involve oscillations within reticulo-thalamo-cortical loops and have been associated with cognitive functions including learning and memory (Fernandez & Luthi, 2020). Spindles are synchronized with other NREM sleep oscillations. Their coupling with cortical slow waves and hippocampus sharp-waves ripples is crucial for triggering reactivation of learning material and promoting hippocampo-neocortical transfer of information in the context of sleep-related memory consolidation (Klinzing, Niethard, & Born, 2019). Such transfer, which requires a fine temporal alignment between these different activities, could be facilitated by respiration (Heck, Kozma, & Kay, 2019). Indeed, respiration was shown to modulate neuronal activity in large-scale subcortical and cortical networks, during both wake (Kluger & Gross, 2021; Zelano et al., 2016) and sleep states (Girin et al., 2021; Tort, Hammer, Zhang, Brankack, & Draguhn, 2021). In the rodent, the respiratory cycle modulates the timing of sharp-wave ripples, whose occurrence probability increases during the early expiration phase (Liu, McAfee, & Heck, 2017). During sleep, respiration promotes the coupling between hippocampal sharp-wave ripples and cortical DOWN/UP state transitions, a neuronal process involved in memory consolidation (Karalis & Sirota, 2022). Behaviourally, the enhancement of recognition memory in humans by nasal breathing strongly supports a role for breathing in the consolidation process (Arshamian, Iravani, Majid, & Lundstrom, 2018). Thus, given the importance of sleep spindles in memory consolidation, and the potential role of respiration in their coupling with other neuronal events, we aimed here at examining whether spindles could be respiration-modulated in humans. Knowing the different functional roles of fast and slow spindles (Fogel & Smith, 2011), we also investigated whether such coupling apply to both spindle types.

## METHODS

### Participants

Twenty healthy subjects (mean ± SD age 31.6 ± 8.3 years; 17 females, 3 males) who had participated in different research projects including a full night polysomnography (PSG) were randomly selected for this study. All participants had benefited from an extended clinical evaluation. Exclusion criteria were: any sleep disorders; any medical or psychiatric conditions; any medication or drug known to influence sleep; shift work; sleep deprivation (ruled out by a 7-day actigraphy prior the PSG). All participants granted informed consent. The study was approved by the Hospices Civils de Lyon Ethics Review Board (N° 22_5757, 10/06/2022).

### Sleep recordings

Night sleep recordings were conducted in the Centre for Sleep Medicine and Respiratory Diseases of Lyon University Hospital between 2017 and 2021. The following signals were recorded with the Deltamed/Natus® acquisition system (sample rate: 256 Hz): EEG (Fp1, Fp2, Fz, C3, C4, Cz, T3, T4, Pz, O1, O2), electro-oculogram, chin and tibialis electromyogram, EKG, nasal airflow (nasal pressure and oronasal thermistor), pulse oximetry, microphone, and respiratory efforts (thoracic and abdominal belts).

### Respiratory recording and processing

Respiratory cycle detection was performed using an algorithm previously described applied on nasal pressure signal (Roux et al., 2006). Briefly, it consists in performing signal smoothing for noise reduction and detection of zero-crossing points, allowing an accurate definition of the inspiration and expiration phases (respectively positive and negative deflections). The inspiration phase starts at the zero-crossing point of the rising phase and ends at the zero-crossing point of the falling phase.

### EEG data analysis

Sleep staging was performed in 30 s epochs using YASA toolbox (Vallat & Walker, 2021), an automated sleep scoring algorithm based on the American Academy of Sleep Medicine recommendations. Only N2 stage epochs were retained for analyses. The same toolbox was used for spindles (12-15 Hz; 0.5-2 s; threshold > 1.5 SD of the root mean square of the sigma-filtered signal) and slow waves (0.2-1.5 Hz, 0.3-1.5 s duration for negative deflection and 0.1-1 s duration for positive deflection, peak-to-peak amplitude 75-350 µV) detection. Fast and slow spindles were separated through a frequency threshold manually set for each participant according to the bimodal distribution of their frequencies. These thresholds ranged from 12.3 to 13.7 Hz (mean: 13.15 Hz). Time-frequency (TF) maps of the NREM sleep signal were obtained using Complex Morlet Wavelet convolutions. Morlet wavelet settings were optimized to capture spindle-related sigma power of the signal as a function of time (frequencies linearly increasing from 10 to 16 Hz in 60 steps and with 20 cycles oscillations, tapering to zero, 10 s duration). TF maps were then sliced into epochs according to onset and offset of each respiratory cycle. TF maps corresponding to each of these epochs were then “stretched” as explained in Roux et al. (2006). Briefly, the time component of these epochs, which can differ from trial to trial, was converted into a phase component defined as [0,2/3Pi] and [2/3Pi, 2Pi] for inspiration and expiration, respectively. As opposed to time representation, the phase representation is common to all epochs. Thus, this phase representation of the respiratory cycle is used as a normalized time basis allowing collecting results in a standardized data format across different participants and providing a way to average oscillatory components of the activity. We then averaged such phase-frequency maps depending on whether they contained spindles or not. Phase-frequency maps containing at least one detected spindle were averaged, allowing spindle-related sigma power representation as shown in figure 1. Spindle starting times and slow wave negative peak times were timestamped thanks to YASA algorithm. These timestamps were converted to phase angles according to the relative time of occurrence during their corresponding breathing cycle.

**Figure 1.**
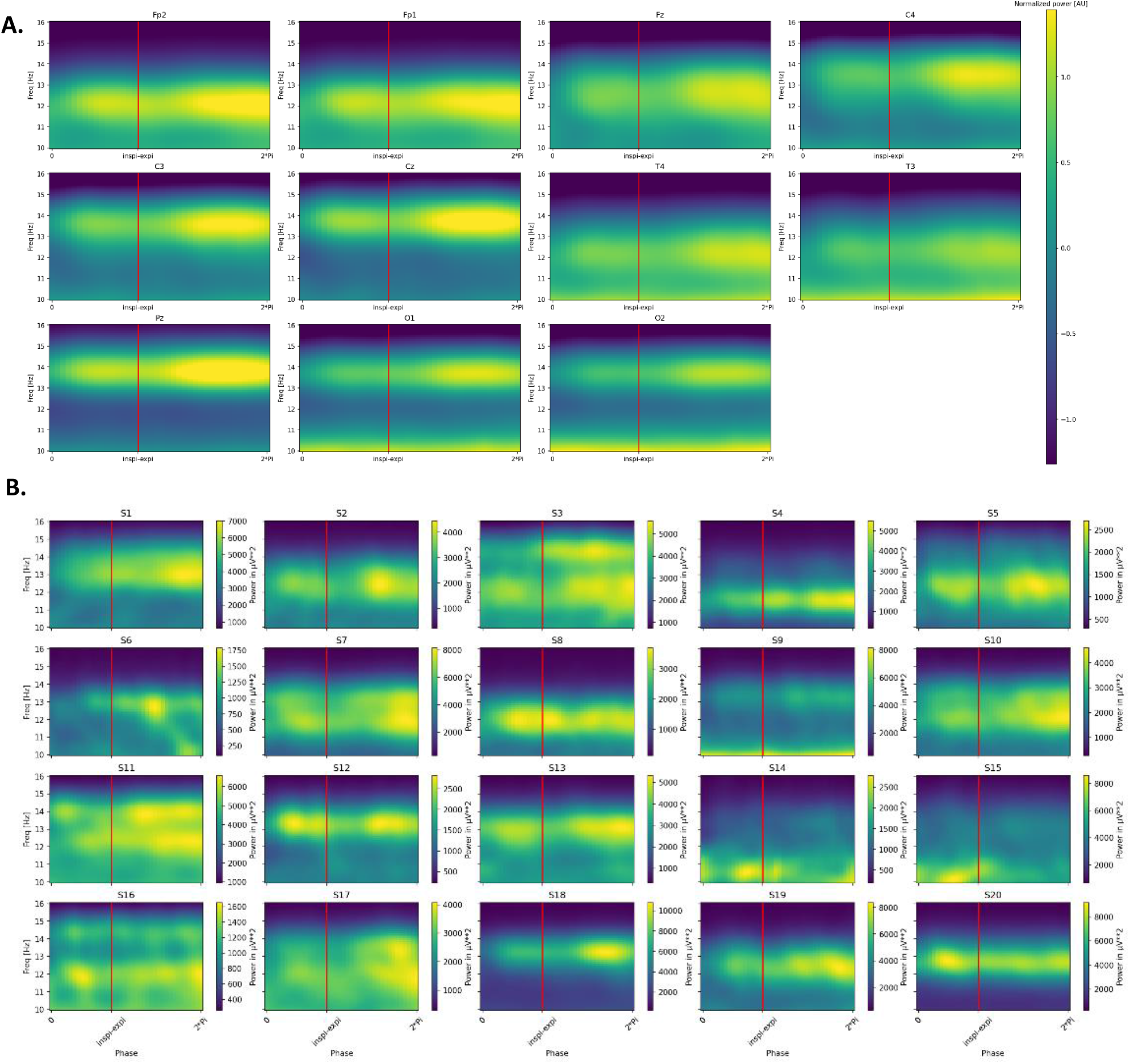
Spindle-related sigma power according to respiration phase. Sigma power is higher during the expiratory phase. Respiration phase is expressed in radians from 0 to 2Pi, beginning from inspiration and ending before the next inspiration. Red vertical line show the timing of the transition from inspiration to expiration. **A:** Mean respiratory phase-frequency map sigma band power (n=20 subjects). Power matrix of each subject and channel has been z-scored before averaging. Note that spindle frequency is lower in anterior (Fp1, Fp2) than in posterior (C3, C4, Cz, Pz) channels. **B:** Individual phase-frequency maps of Fz channel sigma band power. Note the double band representation of the sigma power for some subjects (siuch as S3 and S11), due to the non-overlapping of their slow versus fast spindle-related sigma frequencies.

### Statistical analysis

For each participant, mean direction and vector length of these two sets of phase angles (spindles peaks and slow wave negative peaks) were computed using circular statistics. Rayleigh test was performed to test for non-uniformity of circular distribution of angles and data were considered as non-uniform if p-value returned lower than 0.05.

## RESULTS

### Sleep features

On average 230 ± 31 min of N2 epochs per participant were used for analyses. Sleep parameters are presented in table 1. Considering all channels (n=11 per subject), 5543 ± 2739 spindles were detected per participant (mean density = 2.2 spindles per minute per channel per participant) with a 0.88 ± 0.30 seconds of duration and a 13.4 ± 0.7 Hz frequency, and 2834 ± 1353 slow waves were detected per participant, with a 1.25 ± 0.29 seconds of duration, a 0.84 ± 0.19 Hz frequency, and a negative-to-positive peak mean amplitude of 127 ± 49 µV. Spindle frequency distribution according to channel localization is provided in supplementary Figure 1.

**Table 1.** Sleep parameters during the recorded night. Sleep parameters have been computed for each subject (S1 to S20). Mean and standard deviations (SD) are presented in the last two rows. All units are minutes or percentages (% in columns labels). Abbreviations: Time in Bed (TIB): total duration of the hypnogram ; Total Sleep Period (TSP): duration from first to last epoch of sleep. ; Wake After Sleep Onset (WASO): duration of wake periods within TSP ; Total Sleep Time (TST): total duration of N1 + N2 + N3 + R sleep in TSP ; Sleep Efficiency (SE): TST / TIB * 100 (%) ; Sleep Maintenance Efficiency (SME): TST / TSP * 100 (%) ; W, N1, N2, N3 and R: sleep stages duration. NR = N1 + N2 + N3 ; % (W, … R): sleep stages duration expressed in percentages of TST ; Sleep Onset Latency (SOL): Latency to first epoch of any sleep stage.

| Sub | TIB | TSP | WASO | TST | N1 | N2 | N3 | R | NR | SOL | %N1 | %N2 | %N3 | %R | %NR | %SE | %SME |
| --- | --- | --- | --- | --- | --- | --- | --- | --- | --- | --- | --- | --- | --- | --- | --- | --- | --- |
| S1 | 527,5 | 440 | 32,5 | 407,5 | 30 | 190,5 | 64,5 | 122,5 | 285 | 51,5 | 7,36 | 46,75 | 15,83 | 30,06 | 69,94 | 77,25 | 92,61 |
| S2 | 560,5 | 520 | 45 | 475 | 29,5 | 242,5 | 83 | 120 | 355 | 40,5 | 6,21 | 51,05 | 17,47 | 25,26 | 74,74 | 84,75 | 91,35 |
| S3 | 598,5 | 580,5 | 43,5 | 537 | 36 | 290,5 | 73,5 | 137 | 400 | 18 | 6,7 | 54,1 | 13,69 | 25,51 | 74,49 | 89,72 | 92,51 |
| S4 | 524,5 | 503,5 | 51,5 | 452 | 61 | 241 | 50 | 100 | 352 | 14,5 | 13,5 | 53,32 | 11,06 | 22,12 | 77,88 | 86,18 | 89,77 |
| S5 | 498,5 | 488 | 50 | 438 | 53,5 | 216 | 63,5 | 105 | 333 | 10,5 | 12,21 | 49,32 | 14,5 | 23,97 | 76,03 | 87,86 | 89,75 |
| S6 | 509 | 474 | 44 | 430 | 45,5 | 201 | 109,5 | 74 | 356 | 35 | 10,58 | 46,74 | 25,47 | 17,21 | 82,79 | 84,48 | 90,72 |
| S7 | 523 | 469,5 | 53 | 416,5 | 29,5 | 219,5 | 84 | 83,5 | 333 | 53,5 | 7,08 | 52,7 | 20,17 | 20,05 | 79,95 | 79,64 | 88,71 |
| S8 | 430 | 421,5 | 73,5 | 348 | 40,5 | 177,5 | 45 | 85 | 263 | 8,5 | 11,64 | 51,01 | 12,93 | 24,43 | 75,57 | 80,93 | 82,56 |
| S9 | 542 | 537,5 | 40 | 497,5 | 14,5 | 289 | 72 | 122 | 375,5 | 4,5 | 2,91 | 58,09 | 14,47 | 24,52 | 75,48 | 91,79 | 92,56 |
| S10 | 535 | 483 | 83 | 400 | 31,5 | 220 | 68,5 | 80 | 320 | 32,5 | 7,88 | 55 | 17,12 | 20 | 80 | 74,77 | 82,82 |
| S11 | 430,5 | 417 | 42 | 375 | 13,5 | 185 | 68,5 | 108 | 267 | 13,5 | 3,6 | 49,33 | 18,27 | 28,8 | 71,2 | 87,11 | 89,93 |
| S12 | 539 | 501,5 | 89,5 | 412 | 52 | 214 | 36 | 110 | 302 | 15 | 12,62 | 51,94 | 8,74 | 26,7 | 73,3 | 76,44 | 82,15 |
| S13 | 468,5 | 448,5 | 17,5 | 431 | 23,5 | 220 | 98 | 89,5 | 341,5 | 20 | 5,45 | 51,04 | 22,74 | 20,77 | 79,23 | 92 | 96,1 |
| S14 | 578 | 535 | 29 | 506 | 39,5 | 253,5 | 82 | 131 | 375 | 43 | 7,81 | 50,1 | 16,21 | 25,89 | 74,11 | 87,54 | 94,58 |
| S15 | 528,5 | 517,5 | 32 | 485,5 | 26,5 | 241 | 107,5 | 110,5 | 375 | 11 | 5,46 | 49,64 | 22,14 | 22,76 | 77,24 | 91,86 | 93,82 |
| S16 | 473,5 | 458,5 | 8 | 450,5 | 11 | 232,5 | 95 | 112 | 338,5 | 15 | 2,44 | 51,61 | 21,09 | 24,86 | 75,14 | 95,14 | 98,26 |
| S17 | 548 | 518,5 | 22 | 496,5 | 29,5 | 256,5 | 70,5 | 140 | 356,5 | 21 | 5,94 | 51,66 | 14,2 | 28,2 | 71,8 | 90,6 | 95,76 |
| S18 | 575,5 | 566 | 60,5 | 505,5 | 32 | 254 | 103,5 | 116 | 389,5 | 9 | 6,33 | 50,25 | 20,47 | 22,95 | 77,05 | 87,84 | 89,31 |
| S19 | 574,5 | 558,5 | 64,5 | 494 | 26 | 212,5 | 114 | 141,5 | 352,5 | 16 | 5,26 | 43,02 | 23,08 | 28,64 | 71,36 | 85,99 | 88,45 |
| S20 | 533 | 509,5 | 16,5 | 493 | 29 | 255 | 107,5 | 101,5 | 391,5 | 23,5 | 5,88 | 51,72 | 21,81 | 20,59 | 79,41 | 92,5 | 96,76 |
| Mean | 524,8 | 497,4 | 44,88 | 452,5 | 32,7 | 230,5 | 79,8 | 109,4 | 343,0 | 22,8 | 7,34 | 50,92 | 17,57 | 24,16 | 75,84 | 86,22 | 90,92 |
| SD | 46,13 | 46,44 | 21,85 | 50,03 | 13,14 | 30,99 | 22,6 | 20,05 | 39,41 | 14,71 | 3,2 | 3,2 | 4,46 | 3,41 | 3,41 | 5,8 | 4,58 |

### Respiration features

Participants respiration cycles lasted on average 3.77 ± 0.45 s corresponding to 1,57 ± 0.18 s of inspiration and 2.19 ± 0.3 s of expiration. Detailed mean respiration features computed during N2 sleep stage for each participant are presented in table 1 of supplementary data.

### Spindle-related sigma power coupling with respiration

Spindle related sigma power was extracted using Morlet Wavelet convolutions and computed along respiration cycle phase. As evidenced by the mean respiratory phase-frequency map of sigma band power (Fig.1A), the power was not evenly distributed over the respiratory cycle but was higher during the expiratory phase for all the channels. Individual maps in Fz channel (Fig.1B) revealed that it was true for most of the 20 subjects. Nevertheless, an inter-individual variability appeared in the phase, frequency and/or duration of the episodes of sigma high power.

### Spindles and slow waves events coupling with respiration

When considering the probability of occurrence of spindle events detected on the different channels as a function of the respiratory cycle, Rayleigh test underlined a non-uniform distribution for spindles start times pooled from the twenty subjects (p-value < 0,05). The mean direction indicated a significantly increased probability during the expiration phase (Fig.2A, grey polar distribution for Fz channel). Since NREM spindles have been described as phase-locked to sleep slow waves (Clemens et al., 2007), we explored if slow waves were also respiration-phased. Polar plots of slow waves maxima revealed that they were not as non-uniformly distributed as spindles but still with a significant increase of probability of occurrence around both respiration transition phases (p-value < 0,001) (Fig.2A, green polar distribution for Fz channel). Knowing the different functional role of slow compared to fast spindles, we analysed separately these two spindle populations. We found a significant increase of distribution of spindles toward expiration phase for both slow and fast spindles (Fig.2B dark and light blue presenting slow and fast spindles coupling, respectively). Indeed, mean vector lengths are equal to 0.079 and 0.090 when mean angle are equal to 269°, 264° for the slow and fast spindles, respectively, for Fz channel. Results of other channels are presented in figure 2C and the corresponding circular statistics in supplementary data, table 2.

**Figure 2.**
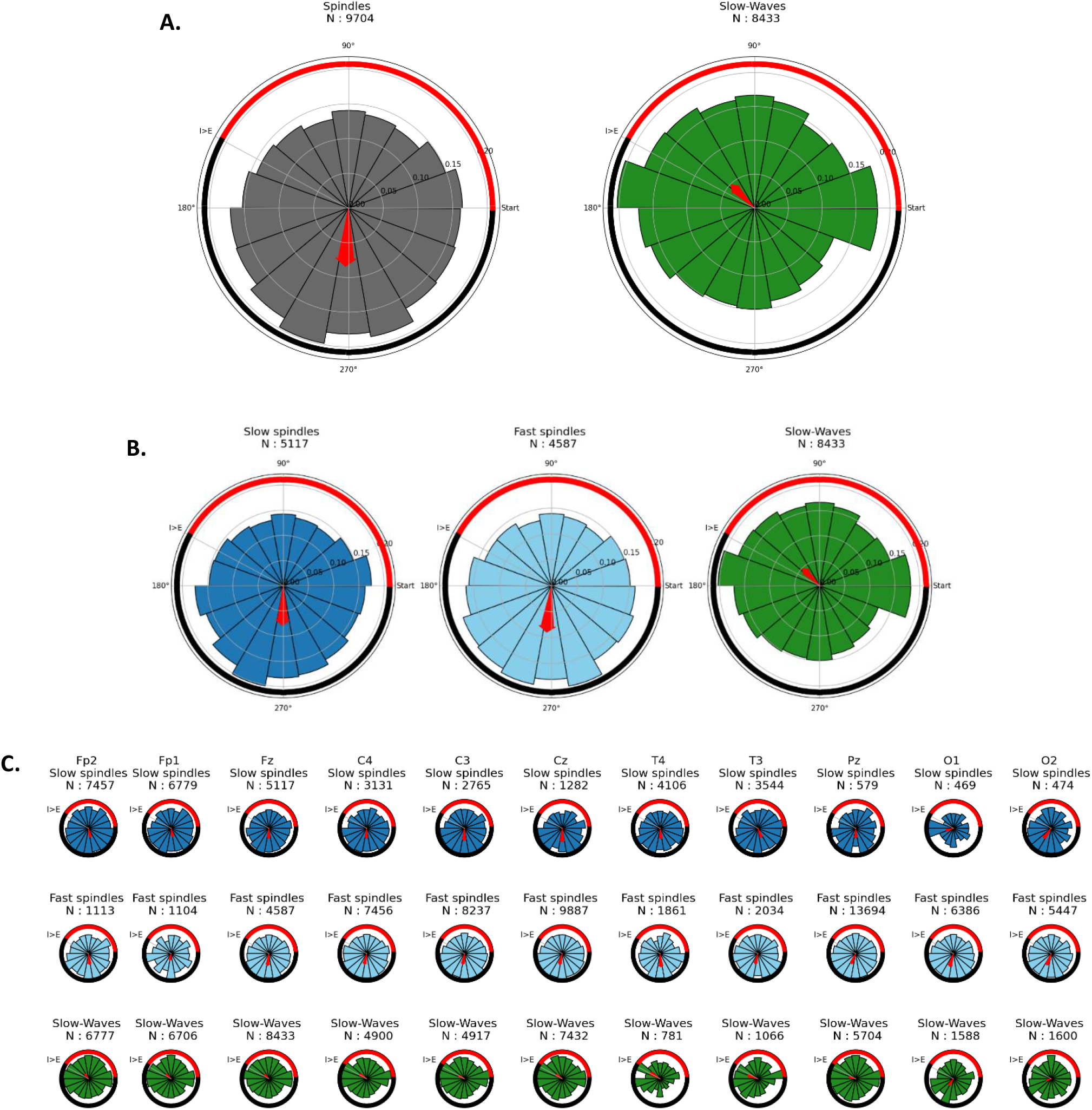
Spindle and slow waves events occurrence according to respiration phase. Spindle start times and slow-wave negative peaks were considered as timing events. Timing events from the 20 subjects were pooled. These events were labelled with the phase angle of occurrence according to breathing cycle and distributed in 18 equal phase bins of respiration (20° per bin). The count of events distributed is presented in title (N). Inspiration and expiration phases are indicated by the red and black lines respectively. A red arrow indicates the mean angular direction, and its length depends on the resultant mean vector length that is represented on the circular ticks. **A**. Polar distribution of spindles (grey) and slow wave (green) events according to respiratory cycle. Mean vector lengths are equal to 0.084 and 0.049 for the spindles and slow-waves polar distributions, respectively. **B**. Polar distribution of slow spindles (dark blue), fast spindles (light blue) and slow wave (green) events according to respiratory cycle. Mean vector lengths are equal to 0.079, 0.090 and 0.049, for the slow spindles, fast spindles and slow-waves polar distributions, respectively. **C**. Polar distribution of slow spindles (dark blue), fast spindles (light blue) and slow wave (green) for each channel events according to respiratory cycle. Circular statistics for each channel are presented in supplementary data, table 2. All distributions presented in A, B and C are significantly non-uniform as assessed by Rayleigh test (p-value < 0.05). Globally, slow and fast spindles occurrence is higher during the expiration phase while slow waves are more distributed near respiration phase transitions (especially inspiration to expiration).

## DISCUSSION

We found a significant coupling between sleep spindles and respiration cycle, with enhanced sigma activity and spindle events (slow or fast) during the second part of the expiration phase. These results show for the first time that sleep oscillations are modulated by breathing in humans.

During wakefulness, local field potential activity in limbic area including the piriform cortex, the amygdala, and the hippocampus, is entrained by nasal breathing, disappearing with oral breathing (Zelano et al., 2016). This suggests that the respiration-locked effect may be driven by the mechano-sensitivity of olfactory sensory neurons to air pressure, with olfactory bulb responses propagating to other brain area (Girin et al., 2021). During sleep however, a recent study in mice revealed that respiratory modulation of sharp wave ripples occurrence and UP/DOWN states is maintained after olfactory deafferentation, suggesting the existence of an ascending respiratory corollary discharge, likely propagating from the brainstem (Karalis & Sirota, 2022). In line with the extensive data showing the persistence of brain responses to various sensory inputs during sleep (Bastuji & Garcia-Larrea, 1999), our results reveal that breathing is able to influence brain activity during N2 sleep in thalamo-cortical networks in humans. The relative contribution of nasal vs ascending respiratory activity should be further investigated.

We observed that slow and fast sleep spindles were locked to the expiratory phase with a high within and between subject reproducibility. This oscillatory pacemaker effect of breathing may enable sleep spindles to synchronize with other brain rhythms including hippocampus sharp wave ripples. Respiration-driven enhanced coupling may facilitate information transfer between distributed brain networks, and play a role in sleep-related memory consolidation process (Karalis & Sirota, 2022). Interestingly, we found a different and smaller breathing effect on slow waves occurrence, suggesting that the oscillation between cortical up and down states may partly depend on respiration phase transitions.

We acknowledge several limitations to the study. First, this was an exploratory study on a small number of participants, with a high proportion of young women (participants were healthy controls selected from a study on idiopathic hypersomnia). Results need to be confirmed in larger cohorts of subjects of various ages with more balanced sex ratio. Second, our study focused on N2 stage as sleep spindles are prominent during this stage. Extending this analysis to N3 sleep stage would enable to further investigate the effect of respiration on slow waves as well as on the well-known coupling between slow waves and spindles.

As a conclusion, respiration phase influences the dynamics of brain activity during N2 sleep. The impact of breathing characteristics on sleep-related cognitive functions, as well as the consequences of sleep-related breathing disorders and mechanical ventilation on this breath-to-brain effect remain to be investigated.

## Data and code sharing statement

The data and code that support the findings of this study are available from the corresponding author upon reasonable request.

## Supplementary data

**Figure 1.**
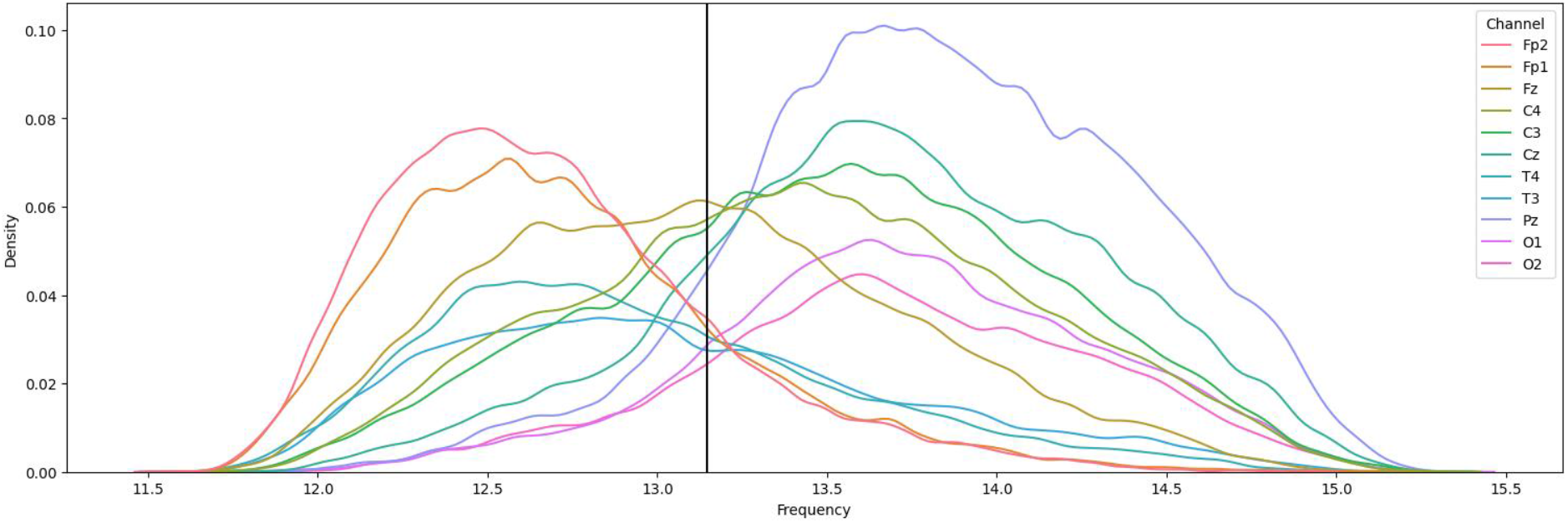
Spindle frequency distribution according to channel localization (N = 10077 ± 3190 spindles per channel pooled from 20 subjects). The bimodal distribution distinguishes between anterior slow spindles (12-13 Hz: Fp1, Fp2, T3, T4) and posterior fast spindles (13-15Hz: Pz, C3, C4, O1, O2). Fz records both slow and fast spindles. Black vertical line represents the mean threshold of separation of spindles into slow and fast (at 13.15Hz).

**Table 1.** Mean respiration features for each subject. Mean respiration features have been computed by averaging features from respiration cycles detected during N2 sleep, for each subject (S1 to S20). Mean and standard deviations (SD) are presented in the last two rows. All units are seconds. Cycle_duration: mean respiration cycle duration ; cycle_ratio: duration of inspiration / cycle_duration ; inspi_duration : duration of inspiration phase, expi_duration : duration of expiration phase.

| Subject | cycle_duration | cycle_ratio | inspi_duration | expi_duration |
| --- | --- | --- | --- | --- |
| S1 | 2,81 | 0,45 | 1,26 | 1,55 |
| S2 | 3,97 | 0,43 | 1,72 | 2,25 |
| S3 | 3,98 | 0,43 | 1,69 | 2,29 |
| S4 | 3,98 | 0,42 | 1,67 | 2,32 |
| S5 | 4,13 | 0,43 | 1,77 | 2,35 |
| S6 | 3,97 | 0,4 | 1,59 | 2,39 |
| S7 | 3,67 | 0,42 | 1,55 | 2,12 |
| S8 | 4,22 | 0,39 | 1,64 | 2,57 |
| S9 | 4,79 | 0,37 | 1,78 | 3,01 |
| S10 | 3,26 | 0,4 | 1,3 | 1,96 |
| S11 | 3,4 | 0,42 | 1,42 | 1,97 |
| S12 | 3,82 | 0,44 | 1,67 | 2,15 |
| S13 | 3,31 | 0,4 | 1,33 | 1,98 |
| S14 | 3,93 | 0,47 | 1,85 | 2,08 |
| S15 | 4,24 | 0,42 | 1,78 | 2,46 |
| S16 | 3,77 | 0,44 | 1,65 | 2,13 |
| S17 | 3,52 | 0,4 | 1,41 | 2,11 |
| S18 | 3,12 | 0,43 | 1,33 | 1,79 |
| S19 | 3,85 | 0,41 | 1,58 | 2,28 |
| S20 | 3,59 | 0,42 | 1,5 | 2,09 |
| Mean | 3,77 | 0,42 | 1,57 | 2,19 |
| SD | 0,45 | 0,02 | 0,18 | 0,3 |

**Table 2.** Circular statistics of distribution of events pooled from 20 subjects, in each channel.

| Channel | Event | N | p-Rayleigh | Rayleigh Significance | Mean Direction | Mean Vector |
| --- | --- | --- | --- | --- | --- | --- |
| Fp2 | Slow spindles | 7457 | < 0.001 | *** | 283 | 0,064 |
| Fp2 | Fast spindles | 1113 | < 0.001 | *** | 276 | 0,083 |
| Fp2 | Slow-Waves | 6777 | < 0.001 | *** | 126 | 0,046 |
| Fp1 | Slow spindles | 6779 | < 0.001 | *** | 282 | 0,063 |
| Fp1 | Fast spindles | 1104 | 0,0232 | * | 260 | 0,058 |
| Fp1 | Slow-Waves | 6706 | < 0.001 | *** | 137 | 0,052 |
| Fz | Slow spindles | 5117 | < 0.001 | *** | 269 | 0,079 |
| Fz | Fast spindles | 4587 | < 0.001 | *** | 264 | 0,09 |
| Fz | Slow-Waves | 8433 | < 0.001 | *** | 135 | 0,049 |
| C4 | Slow spindles | 3131 | < 0.001 | *** | 268 | 0,074 |
| C4 | Fast spindles | 7456 | < 0.001 | *** | 258 | 0,081 |
| C4 | Slow-Waves | 4900 | < 0.001 | *** | 147 | 0,067 |
| C3 | Slow spindles | 2765 | < 0.001 | *** | 268 | 0,088 |
| C3 | Fast spindles | 8237 | < 0.001 | *** | 261 | 0,079 |
| C3 | Slow-Waves | 4917 | < 0.001 | *** | 152 | 0,053 |
| Cz | Slow spindles | 1282 | < 0.001 | *** | 270 | 0,104 |
| Cz | Fast spindles | 9887 | < 0.001 | *** | 256 | 0,081 |
| Cz | Slow-Waves | 7432 | < 0.001 | *** | 147 | 0,059 |
| T4 | Slow spindles | 4106 | < 0.001 | *** | 275 | 0,081 |
| T4 | Fast spindles | 1861 | < 0.001 | *** | 274 | 0,102 |
| T4 | Slow-Waves | 781 | < 0.001 | *** | 149 | 0,111 |
| T3 | Slow spindles | 3544 | < 0.001 | *** | 292 | 0,071 |
| T3 | Fast spindles | 2034 | < 0.001 | *** | 256 | 0,078 |
| T3 | Slow-Waves | 1066 | < 0.001 | *** | 160 | 0,09 |
| Pz | Slow spindles | 579 | 0,0371 | * | 271 | 0,075 |
| Pz | Fast spindles | 1369 | < 0.001 | *** | 247 | 0,08 |
| Pz | Slow-Waves | 5704 | < 0.001 | *** | 164 | 0,051 |
| O1 | Slow spindles | 469 | 0,053 | ns | 196 | 0,079 |
| O1 | Fast spindles | 6386 | < 0.001 | *** | 254 | 0,099 |
| O1 | Slow-Waves | 1588 | < 0.001 | *** | 235 | 0,068 |
| O2 | Slow spindles | 474 | 0,0147 | * | 233 | 0,094 |
| O2 | Fast spindles | 5447 | < 0.001 | *** | 249 | 0,093 |
| O2 | Slow-Waves | 1600 | 0,0785 | ns | 198 | 0,04 |

